# Telomere-to-telomere, accurate, and gapless genome assembly (TTAGGA) of the Korean Jindo dog with a single-contig Y chromosome

**DOI:** 10.64898/2026.05.17.725804

**Authors:** Hyoungjin Choi, Jong-Seok Kim, Yoonsung Kwon, Sangsoo Park, Sungwon Jeon, Jihun Bhak, Donghyun Shin, Yookyung Choi, Kyungwhan An, Doug-Young Ryu, Woon Kee Paek, Daeui Park, Jaemin Kim, Mikkel-Holger S. Sinding, Youngjae Choe, Bo-Ra Hyun, Sung-kyung Lee, Jong Bhak

## Abstract

Complete canine reference genomes are essential for studies of structural variation, sex-chromosome evolution, and breed-specific architecture, yet existing references remain incomplete on the Y chromosome and retain internal gaps across multiple chromosomes. Here, we introduce TTAGGA (Telomere-to-Telomere, Accurate, Gapless Genome Assembly), a stricter completeness standard requiring chromosomes assembled end-to-end with zero internal gaps and Merqury consensus QV ≥ 50, and present Jindo1-G-TTAGGA, the first canine assembly to meet this standard, achieved under a stringent QV ≥ 60 threshold. From 813 Gb (∼340× coverage) of multi-platform data (PacBio HiFi ∼150×, ONT ultra-long ∼103×, parental Illumina ∼40× per parent) generated from a male Korean Jindo dog, trio binning with hifiasm produced two haplotype-resolved assemblies of 2,441.6 Mb (Hap1, maternal, chrX-carrying) and 2,340.5 Mb (Hap2, paternal, chrY-carrying). Both haplotypes are gap-free at every internal position, with canonical telomeric repeats verified at both ends of all 39 chromosomes (tidk), Merqury consensus QV of 78.0 (Hap1) and 76.8 (Hap2), trio switch-error rates below 0.13%, BUSCO completeness of 99.3% (Hap1) and 96.4% (Hap2; the lower value reflects absent X-linked orthologues in the Y-bearing haplotype), and Genome Continuity Index values of 98.2 (Hap1) and 94.7 (Hap2). Hap2 carries a single 21,255,890 bp gap-free Y chromosome with TTAGGG telomeric repeats at the q-arm terminus and an acrocentric satellite-rich p-arm, representing a 5.4-fold increase over the 3.94 Mb chrY of ROS_Cfam_1.0 and adding approximately 14 Mb of newly resolved Y-linked sequence; this corresponds to roughly 79% of the cytogenetically estimated 27 Mb full-length canine chrY. Jindo1-G-TTAGGA provides a chromosome-scale, haplotype-resolved, gap-free canine reference for studies of canine structural variation, sex-chromosome evolution, and canid phylogenomics.

## Introduction

Complete reference genomes are essential for interpreting genetic variation, tracing evolutionary histories, and enabling precision medicine and selective breeding programmes^1-3^. In domestic dogs (Canis lupus familiaris), which share a deep coevolutionary relationship with humans and serve as critical models for genetic and biomedical research^4-6^, high-quality genome references are key to understanding the genetic basis of phenotypic diversity, disease susceptibility, and behavioural traits^7,8^.

The first canine reference genome, CanFam1.0, was published in 2005 from a female Boxer named Tasha^9^, and subsequent versions have been derived predominantly from female individuals (CanFam3.1, UU_Cfam_GSD_1.0 from a female German Shepherd Dog^10^, CanFam6, and CanFam_GSD-Nala^11^). The male-derived references that do include chrY (Wags^12^, the Schall 2023 Bernese and Cairn Terrier assemblies^13^, ROS_Cfam_1.0, and Yella v2 (GCA_031165255.1)^14^) have introduced incremental improvements with Sanger, PacBio, and ONT long-read technologies but remain fragmented, with unresolved internal gaps, incomplete repetitive regions, and missing subtelomeric sequence. The Y chromosome is particularly poorly resolved across all publicly available male canine references. The most contiguous prior chrY assembly comes from ROS_Cfam_1.0 (GenBank GCA_014441545.1)^15^, which provides only 3.94 Mb of chrY at chromosome scale together with ∼2.37 Mb of unplaced chrY-assigned scaffolds, approximately 23% of the cytogenetically expected ∼27 Mb full-length canine chrY^16^. Other male references recover chrY only as fragmented multi-scaffold sequence (Wags, Yella v2, the Schall 2023 assemblies; see Table 3) or lack chrY entirely. The Y chromosome harbours genes involved in male fertility, sex determination, and paternal lineage evolution^17^, but no canid has yet been represented by a chromosome-scale, gap-free Y chromosome.

The recently released Beagle near-T2T canine assembly^18^ (GCA_044048985.1) reaches telomere-to-telomere completeness for 29 of 39 chromosomes, with 13 internal gaps reported in the deposited metadata. A haplotype-resolved, chrY-inclusive canine reference combining these features with zero internal gaps has not yet been described.

The Korean Jindo dog, designated as Natural Monument No. 53 of Korea^19^, is recognised as a basal East Asian breed^20-22^. As with other indigenous lineages, it has experienced limited recent admixture with modern Western breeds, making it a useful reference point for East Asian breeds that remain underrepresented in canine genomics. A complete, male-derived, haplotype-resolved, and gapless canine reference had not previously been generated from the Jindo lineage, or from any East Asian breed.

Recent breakthroughs in long-read sequencing and assembly have made it possible to reconstruct entire chromosomes from one telomere to the other, including highly repetitive centromeric and heterochromatic regions^23-25^. We introduce TTAGGA (Telomere-to-Telomere, Accurate, Gapless Genome Assembly) to denote a stricter standard in which every chromosome reaches both telomeres and contains zero internal gaps. For the present work, we applied this definition with a stringent threshold of Merqury consensus QV ≥ 60, comparable to the consensus accuracy of the T2T-CHM13 human reference (Nurk et al. 2022)^23^. Applying an integrated multi-platform strategy that combines Oxford Nanopore Technologies ultra-long reads^26,27^ and PacBio HiFi long reads^28,29^ together with parental Illumina short reads for trio binning, we generated the first haplotype-resolved, chrY-inclusive, gap-free canine TTAGGA assembly from a male Korean Jindo dog. This assembly resolves all 39 chromosomes on both haplotypes, including a single-contig 21.26 Mb Y chromosome, a 5.4-fold increase over the 3.94 Mb chrY of ROS_Cfam_1.0. Jindo1-G-TTAGGA thus provides a haplotype-resolved, chrY-inclusive canine reference with zero internal gaps on every chromosome, and establishes a foundation for chromosome-scale analyses of canine sex-chromosome architecture, structural variation, repeat content, and paternal-lineage genetics.

## Results

### Sequencing Data Generation and Quality Control

We generated a total of 813 Gb of sequencing data (∼340× genome coverage) for a male Korean Jindo dog (Baeksan, meaning white mountain), together with parental short-read data for trio binning (Table 1; Supplementary Figure S1). PacBio HiFi sequencing on the Revio platform produced 349.9 Gb (∼150× coverage) across six flowcells, with mean read length 18-19 kb and mean QV > 29. After quality filtering (read length ≥ 1 kb, QV ≥ 20, with 150 bp end-trimming), 317.6 Gb (17.3 million reads) were retained.

**Table 1.** Sequencing summary of the three data types used for Jindo1-G-TTAGGA assembly.

| Technology | Platform / Flowcell | Total yield (Gb) | Coverage (×) | N50 / Read quality |
| --- | --- | --- | --- | --- |
| PacBio HiFi | Revio (6 flowcells) | 349.9 | $\sim 150$ | 18-19 kb; QV $> 29$ |
| ONT ultra-long | PromethION P2 Solo | 258.8 | $\sim 103$ | N50 37.8 kb |
| Illumina (trio) | NovaSeq 6000 | $\sim 250$ (total) | 39 (dam), 42 (sire) | QV $> 30$ |

ONT ultra-long sequencing on the PromethION P2 Solo platform produced 258.8 Gb (∼103× coverage) with N50 of 37.8 kb. Reads were filtered to length ≥ 50 kb and QV ≥ 15 with 2 kb end-trimming, yielding 76.8 Gb (∼30×; N50 101.3 kb) for assembly.

For trio binning, Illumina short-read sequencing was performed for both parents on the NovaSeq 6000 platform, yielding 92 Gb (39× coverage) for the dam and 105 Gb (42× coverage) for the sire after adapter trimming. Parental k-mer separation was performed with yak; haplotype 1 (Hap1, maternally inherited) was assigned to the dam-derived haplotype (carrying chrX in this male individual), and haplotype 2 (Hap2, paternally inherited) was assigned to the sire-derived haplotype (carrying chrY). HiFi-based k-mer plots resolved heterozygous and homozygous peaks at 68× coverage.

### Haplotype-resolved TTAGGA Assembly

Initial diploid assembly was performed with hifiasm v0.25.0-r726^30^ using trio binning, integrating quality-filtered HiFi reads (∼127×), high-quality ONT ultra-long reads (∼30×), and parental-specific yak k-mer databases. The initial diploid output comprised 81 scaffolds for Hap1 (dam-derived; 2,441.6 Mb; scaffold N50 65.4 Mb; 4 residual gaps) and 71 scaffolds for Hap2 (sire-derived; 2,340.5 Mb; scaffold N50 65.4 Mb; 2 residual gaps). Hap1 contained the X chromosome (126 Mb) and Hap2 contained the Y chromosome, consistent with the trio-binning assignment. After scaffolding, ten rounds of gap closing and seven rounds of polishing (Methods), all residual gaps were eliminated on both haplotypes. The final assemblies satisfy the TTAGGA criteria, with Merqury consensus QV of 78.0 (Hap1) and 76.8 (Hap2) and switch-error rates below 0.13%. After all assembly steps, the final Hap2 Y chromosome was resolved as a single 21,255,890 bp (21.26 Mb) gap-free contig (described in detail in the chrY section below).

Telomere detection with tidk^31^ (≥ 100 TTAGGG/CCCTAA repeats within 5 kb of scaffold ends) confirmed telomeric repeats at both ends of all 39 chromosomes in both haplotypes. BUSCO v6.0.0 (carnivora_odb12) genome-mode completeness reached 99.3% (Hap1) and 96.4% (Hap2), and the Genome Continuity Index (GCI) reached 98.2 (Hap1) and 94.7 (Hap2), exceeding all publicly available canine reference assemblies evaluated under the same lineage database.

### Comparison with Existing Canine Reference Assemblies

The overall assembly workflow combining trio binning, hifiasm-based diploid assembly, gap closing, and polishing is summarised in Figure 1A, with the k-mer multiplicity spectrum of the proband Jindo individual shown in Figure 1B. Among all publicly available canine reference assemblies, Jindo1-G-TTAGGA combines (i) two fully assembled haplotypes, (ii) chromosome-scale inclusion of the Y chromosome, and (iii) zero internal gaps on both haplotypes (Table 2). Trio-binning-based parental assignment provides an additional advantage for sex-chromosome work but is not in itself a defining TTAGGA property. The Sanger-era references CanFam3.1 and UU_Cfam_GSD_1.0 contain on the order of 10^3^-10^4^ internal gaps, and long-read PacBio Sequel-based references (CanFam6, ROS_Cfam_1.0) carry several hundred. Jindo1-G-TTAGGA is also the first canine reference with a single-contig Y chromosome (21.26 Mb), a 5.4-fold increase over the longest previously assembled canine chrY.

**Table 2.** Comparison of Jindo1-G-TTAGGA with publicly available canine reference assemblies. Whole-genome internal gap counts and chrY lengths were measured directly from each reference FASTA. For per-chrY gap counts and N base totals, see Table 3.

| Reference | Breed | Sex | Technology | Assembly level | chrY (Mb) | Gaps |
| --- | --- | --- | --- | --- | --- | --- |
| **Jindo1-G-TTAGGA (this study)** | **Korean Jindo** | **Male** | **ONT + HiFi + Trio short** | **TTAGGA (gap-free)** | **21.26** | **0** |
| Beagle near-T2T 2025 | Beagle | Female | ONT + HiFi | Near-T2T | — | 13 |
| ROS Cfam 1.0 | Labrador | Male | PacBio Sequel | Chromosome | 3.94 | 951 |
| Wags (Edwards 2021) | Basenji | Male | PacBio + Hi-C | Chromosome | 3.63 | 6 |
| CanFam6 (Tasha) | Boxer | Female | PacBio Sequel | Chromosome | — | 1,200 |
| CanFam_GSD-Nala (Field 2020) | German Shepherd | Female | PacBio + Bionano | Chromosome | — | 700 |
| UU_Cfam_GSD_1.0 | German Shepherd | Female | Sanger + PacBio | Chromosome | — | 15,000 |
| CanFam3.1 | Boxer | Female | Sanger | Chromosome | — | 23,800 |

**Figure 1.**
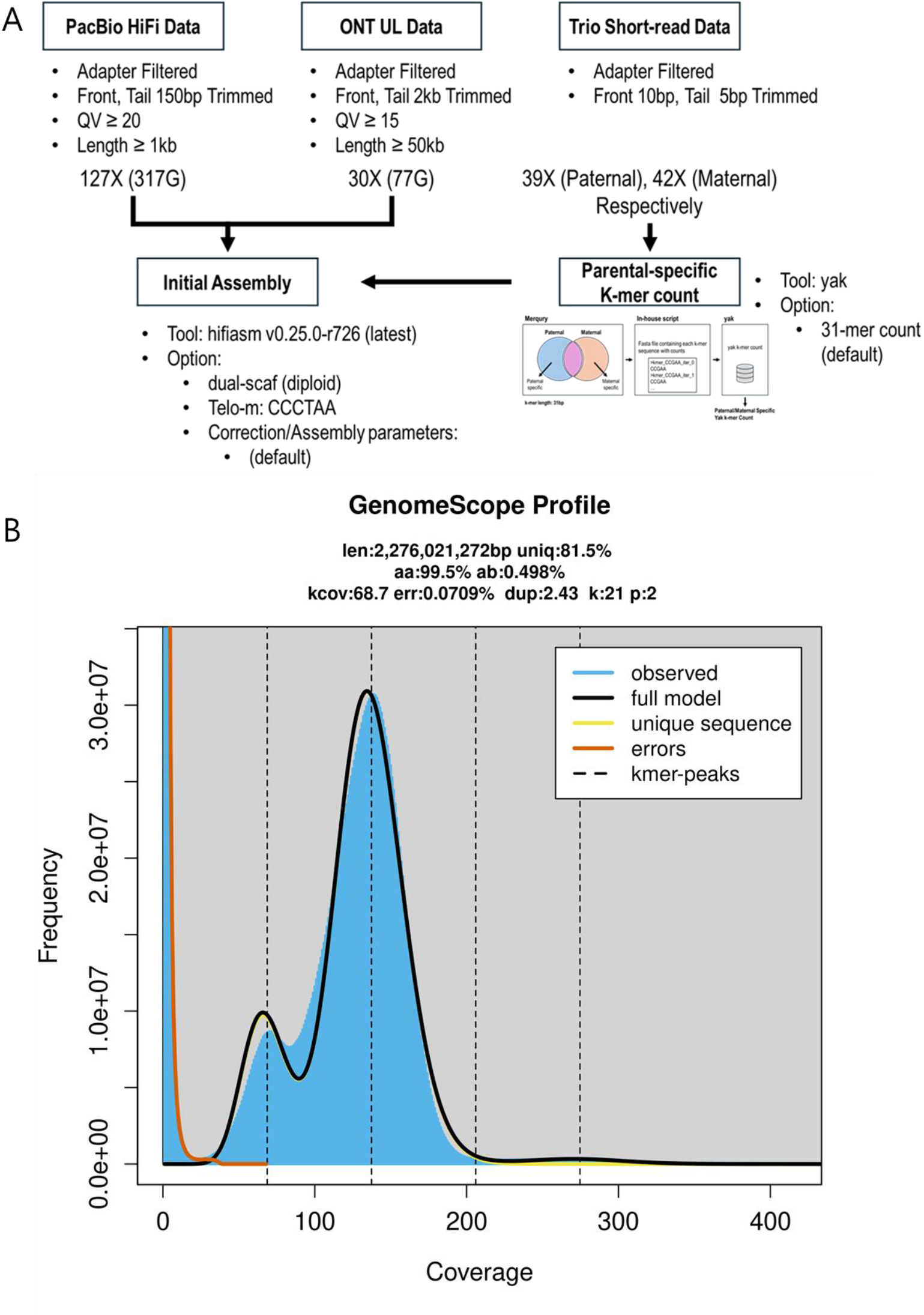
Genome assembly overview of Jindo1-G-TTAGGA. (A) Assembly workflow combining trio binning of parental short reads with hifiasm-based diploid assembly using PacBio HiFi and ONT ultra-long reads, followed by ten rounds of gap closing and seven rounds of polishing. (B) K-mer multiplicity spectrum of the proband Jindo dog (k = 21) generated with GenomeScope2.

### Whole-Genome Synteny against ROS_Cfam_1.0

Whole-genome synteny against ROS_Cfam_1.0 (Labrador Retriever; GCA_014441545.1) was generated using MUMmer (nucmer + delta-filter) for both haplotypes, and structural variants were categorised with SyRI^32^. ROS_Cfam_1.0 was chosen as the primary synteny reference because it contains the most contiguous chromosome-scale chrY (3.94 Mb; Table 3) among publicly available male canine references. Other male references (Wags, Yella v2, the Schall 2023 assemblies) recover chrY only as fragmented chromosome-scale and unplaced scaffolds.

**Table 3.** Y chromosome status across all eight publicly available male canine genome assemblies plus Jindo1-G-TTAGGA Hap2. All chrY lengths, internal gap counts, and total N base counts in chrY scaffolds were measured directly from each reference FASTA. Internal gaps are defined as N stretches ≥ 100 bp at non-terminal positions (> 1 kb from scaffold ends). The “chrY chr-scale (Mb)” column shows the length of the chromosome-scale chrY scaffold (or “(no chr-scale)” if the assembly contains no chromosome-scale chrY); the “Unplaced (Mb)” column shows total length of chrY-assigned unplaced contigs. The “Gaps (N bases)” column shows internal gap count followed by total N bases in parentheses across both chromosome-scale and unplaced chrY scaffolds.

| Assembly | Breed | Technology / Year | chrY chr-scale (Mb) | Unplaced (Mb) | Gaps (N bases) |
| --- | --- | --- | --- | --- | --- |
| **Jindo1-G-TTAGGA Hap2 (this study)** | **Korean Jindo** | **ONT + HiFi + Trio (2026)** | **21.26** | **0** | **0** |
| ROS_Cfam_1.0 | Labrador (SID07034) | PacBio Sequel (2020) | 3.94 | 2.37 | 7 (687,168) |
| Wags | Basenji (male; GCA_004886185.2) | PacBio + Hi-C (2021) | 3.63 | 0 | 6 (600) |
| Cairn Terrier CA611 | Cairn Terrier | ONT (Schall 2023) | 3.54 | 2.14 | 13 (1,300) |
| Bernese Mountain Dog OD | Bernese | ONT (Schall 2023) | 3.31 | 1.96 | 18 (1,800) |
| Yella v2 (Labrador) | Labrador | ONT (GCA_031165255.1) | — (no chr-scale) | 5.37 (3 contigs) | 14 (1,400) |
| Yella alternate haplotype | Labrador | ONT (GCA_012044875.1) | absent | — | — |
| Yella principal haplotype | Labrador | ONT (GCA_012045015.1) | absent | — | — |
| Whippet 2024 | Whippet | PacBio HiFi Revio (2024) | absent | — | — |

Both Jindo haplotypes are largely collinear with ROS_Cfam_1.0 across all 38 autosomes, with focal divergences localised to subtelomeric, centromeric, and chrY regions (Figure 2). Hap1 (maternally inherited, dam-derived) contains 255 syntenic blocks spanning 2.32 Gb, together with 116 inversions, 41 translocations, 3.84 million SNPs, 854,000 insertions, and 673,000 deletions relative to ROS_Cfam_1.0. Hap2 (paternally inherited, sire-derived) shows a structurally similar profile: 215 syntenic blocks (2.20 Gb), 82 inversions, 35 translocations, 3.73 million SNPs, 828,000 insertions, and 648,000 deletions. The dominance of syntenic blocks over rearranged regions indicates broad structural conservation between the Jindo and Labrador lineages. Full structural-variant categorisation across all variant classes is reported as Supplementary Table S5.

**Figure 2.**
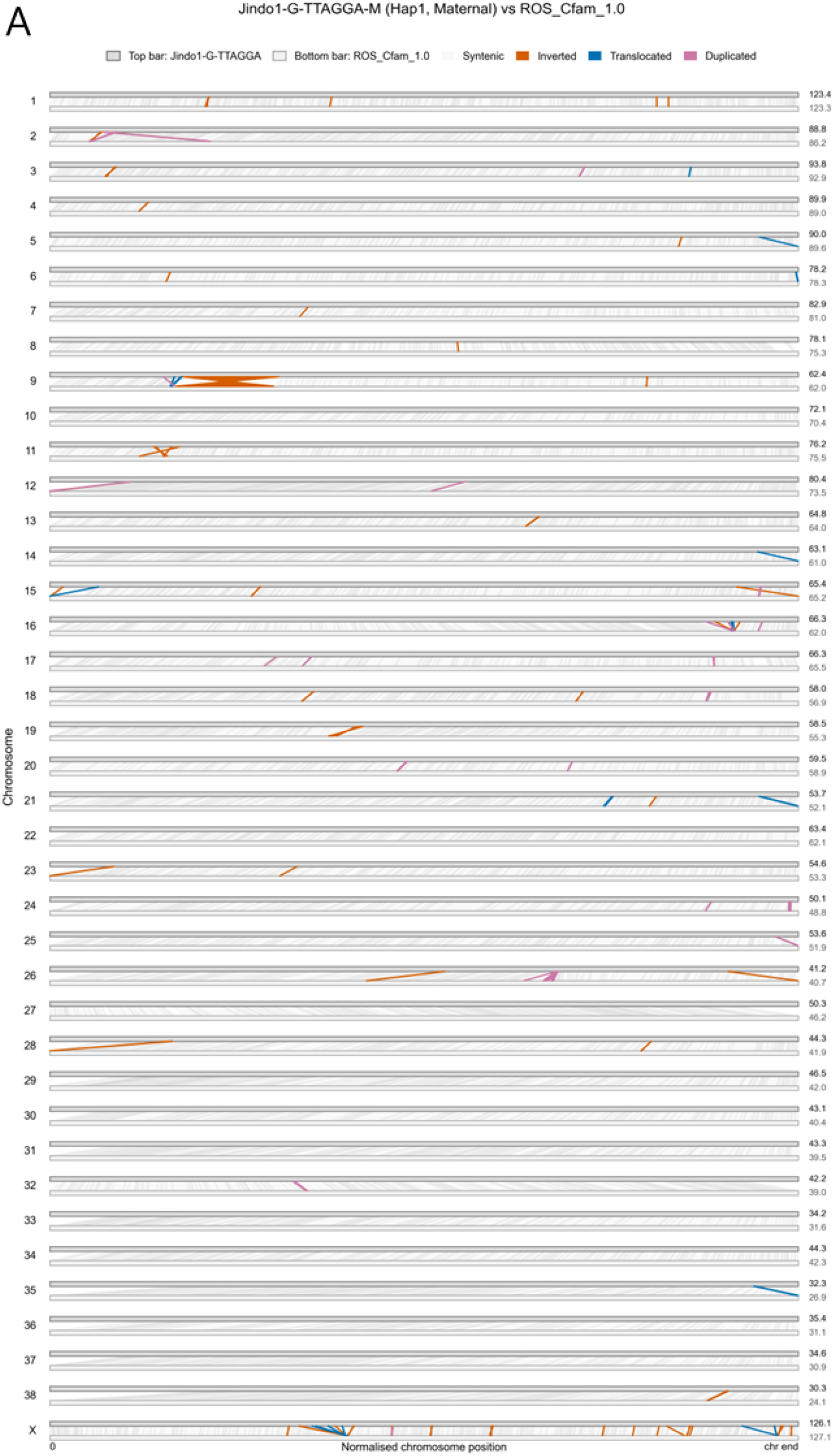
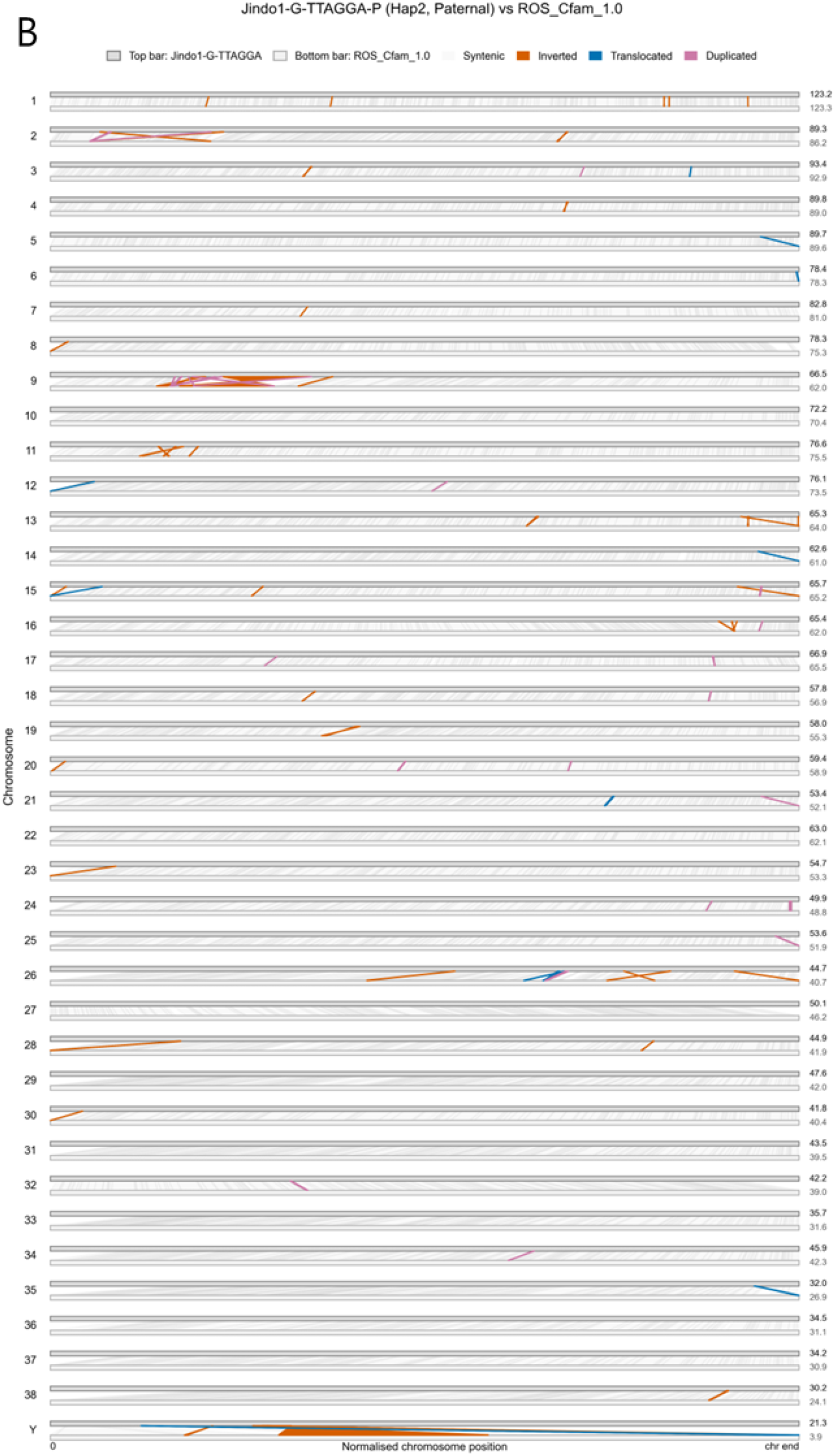
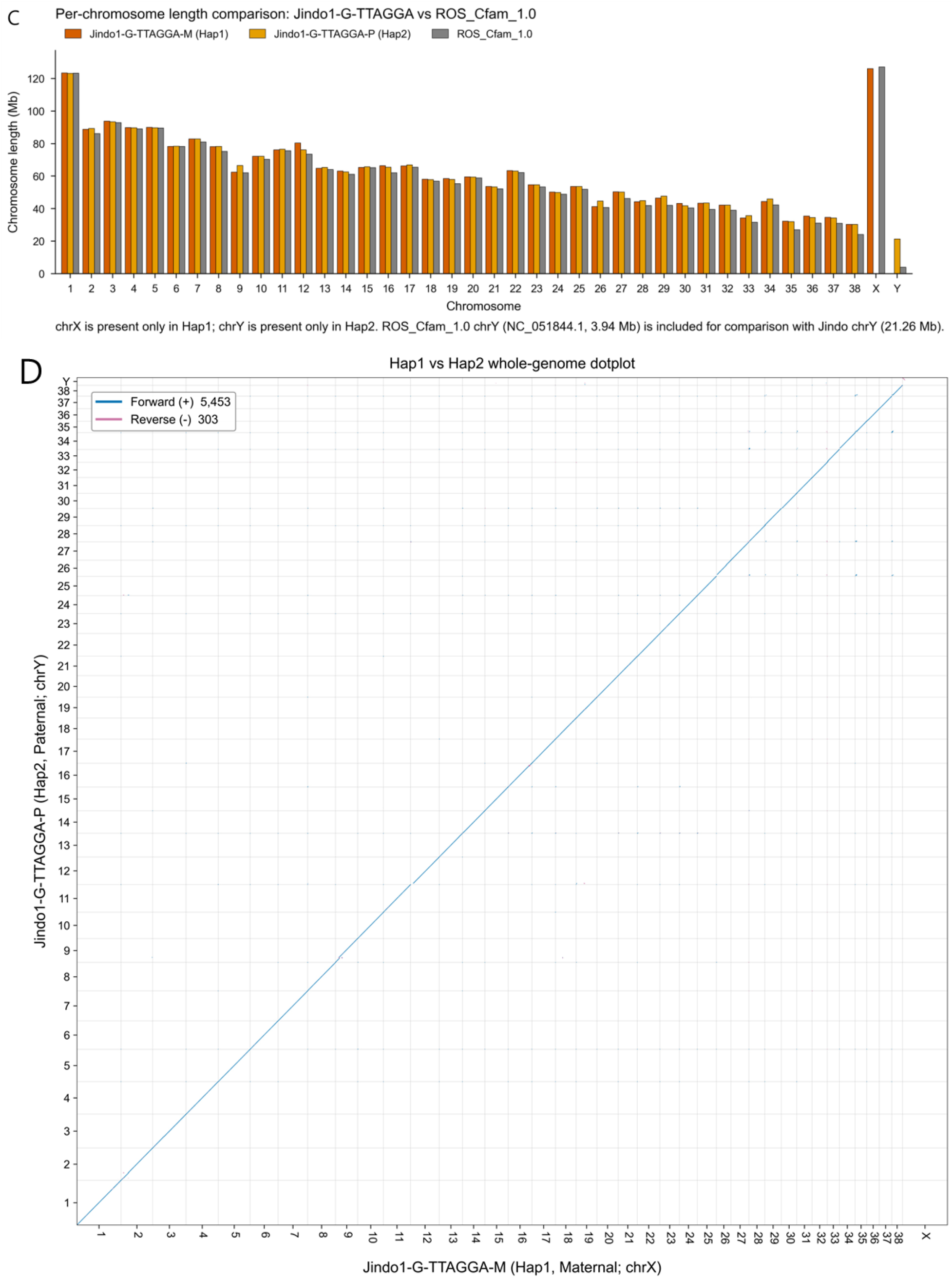
Comparative synteny and chromosome architecture between Jindo1-G-TTAGGA and ROS_Cfam_1.0. (A, B) Whole-genome TSAR plots (TTAGGA Synteny And Rearrangement plots) of (A) Hap1 (Jindo1-G-TTAGGA-M; maternally inherited; chrX-carrying) and (B) Hap2 (Jindo1-G-TTAGGA-P; paternally inherited; chrY-carrying) against ROS_Cfam_1.0. Each row pairs a Jindo chromosome (top, dark grey bar) with the orthologous ROS_Cfam_1.0 chromosome (bottom, light grey bar); chromosome bars are length-normalised. Ribbons drawn between the bars represent alignment blocks ≥ 10 kb and are colour-coded as syntenic (pale grey), inverted (red), translocated (blue), or duplicated (pink); the colour palette follows the Wong (2011) colourblind-safe set. chrX appears only in panel A and chrY only in panel B. (C) Per-chromosome length comparison for all 38 autosomes plus chrX and chrY, with three side-by-side bars per chromosome representing Hap1 (red), Hap2 (yellow), and ROS_Cfam_1.0 (grey). chrX is present only in Hap1 and chrY only in Hap2; ROS_Cfam_1.0 chrY (NC_051844.1, 3.94 Mb) is shown for comparison with the Jindo Hap2 chrY (21.26 Mb). Numerical values are tabulated in Supplementary Table S3. (D) Whole-genome dotplot of Hap2 (y-axis) against Hap1 (x-axis) from alignment blocks ≥ 10 kb, with forward-strand blocks in blue and reverse-strand blocks in pink.

The largest single inversion detected in the synteny analysis was an 8 Mb region on chromosome 9 (Hap1: chr9:11,050,758-19,079,402; Hap2: chr9:15,287,232-23,012,333), present concordantly in both Jindo haplotypes. This same chr9 region has previously been reported as an orientation artefact in earlier canine references: discordant orientations were noted between CanFam3.1 (Boxer Tasha) and the long-read assemblies CanFam_GSD-Nala (Field et al. 2020) and CanFam_Bas (Edwards et al. 2021), and Edwards et al. (2021) attributed the signal to a reference-specific assembly artefact rather than a true biological inversion. Direct chr9-versus-chr9 alignment between Jindo Hap1 and the Beagle near-T2T 2025 assembly shows the same overall orientation. The 8 Mb signal observed in our Jindo-versus-ROS_Cfam_1.0 alignment is therefore attributable to an inherited orientation artefact in the ROS_Cfam_1.0 assembly, not to a Jindo-specific inversion.

Hap1 chrX is largely collinear with ROS_Cfam_1.0 chrX, with inverted regions accounting for less than 0.6% of total chrX length. This indicates that the two assemblies are structurally conserved at chrX and that the bulk of male-female architectural differences in this assembly are restricted to the Y chromosome. Hap2 carries chrY rather than a second X and was therefore not included in this chrX comparison. Comparison of Hap2 chrY against the ROS_Cfam_1.0 chrY scaffold (3.94 Mb) recovered 1 inversion (121 kb), 9 inverted translocations, 10,437 SNPs, 2,246 insertions, and 1,480 deletions across the 4 Mb of mappable overlap. The remaining 14 Mb of the Jindo Hap2 chrY has no counterpart in ROS_Cfam_1.0 and corresponds to newly resolved Y-linked sequence (see chrY section and Figure 3).

**Figure 3.**
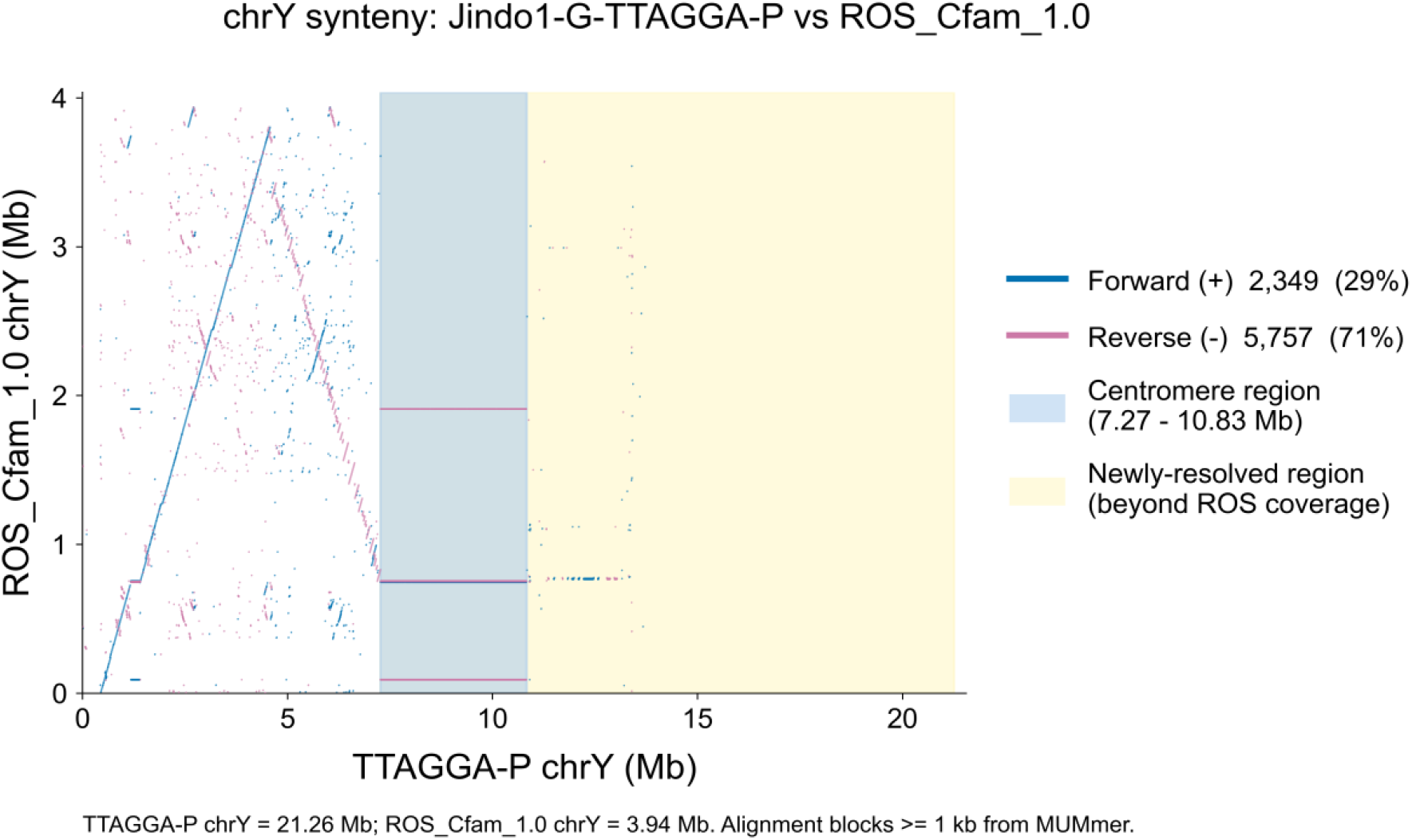
chrY synteny dotplot of Jindo1-G-TTAGGA-P against ROS_Cfam_1.0. Dotplot of the 21.26 Mb gap-free Jindo1-G-TTAGGA-P chrY (Hap2, paternal haplotype; x-axis) against the 3.94 Mb chrY scaffold of ROS_Cfam_1.0 (NC_051844.1; y-axis). Alignments were generated with MUMmer and filtered to blocks ≥ 1 kb. The blue-shaded interval (7.27-10.83 Mb on Jindo chrY) marks the centromere region. The yellow-shaded interval (7.27-21.26 Mb) marks the region with no counterpart in ROS_Cfam_1.0.

### Single-Contig Gap-Free Y Chromosome (21.26 Mb)

The Y chromosome posed a particular challenge because the most complete previously available canine chrY (ROS_Cfam_1.0) recovers only 3.94 Mb at chromosome scale (approximately 15% of the cytogenetically expected ∼27 Mb full-length canine chrY when considering chromosome-scale sequence alone, or ∼23% when 2.37 Mb of unplaced chrY-assigned scaffolds are also counted)^16^. Because reference-based scaffolding cannot exploit telomeric anchors that are absent from existing canine references, we employed a local assembly strategy. Sire-binned reads (HiFi and ONT, i.e., reads assigned to Hap2) were aligned to the dam-derived Hap1 assembly with Winnowmap2; reads mapping to regions lacking telomeric ends were extracted and locally assembled with hifiasm in haploid mode. The resulting assembly was resolved into a single 21,255,890 bp gap-free contig spanning the entire Y chromosome through iterative gap closing and polishing (Methods).

The Hap2 chrY of Jindo1-G-TTAGGA spans 21,255,890 bp in a single gap-free contig with TTAGGG telomeric repeats at the q-arm terminus (53 repeats within the last 10 kb) and an acrocentric satellite-rich p-arm consistent with the heterochromatic short arm characteristic of mammalian acrocentric Y chromosomes. Direct measurement across the eight publicly available male canine assemblies shows that all of them are incomplete: three of the eight (Yella alternate, Yella principal, Whippet 2024 PacBio HiFi Revio) contain no chrY sequence at all, and the remaining five recover only 3.31-3.94 Mb at chromosome scale with 6-18 unresolved internal gaps (Table 3). Yella v2 lacks a chromosome-scale chrY scaffold entirely and carries 5.37 Mb across three unplaced contigs. The 21.26 Mb gap-free Jindo Hap2 chrY thus represents more than a five-fold increase over the longest previously assembled canine chrY.

Synteny alignment of Jindo Hap2 chrY against the ROS_Cfam_1.0 chrY scaffold (MUMmer/nucmer asm5; alignment blocks ≥ 1 kb; Figure 3) shows that the entire 3.94 Mb of ROS_Cfam_1.0 chrY maps to the proximal arm of the Jindo chrY, with no alignment extending into the distal 14 Mb. This 14 Mb (Jindo chrY positions ∼7-21 Mb) represents previously unresolved Y-linked sequence and brings the canine Y chromosome to a length at which palindromic and ampliconic architecture, paternal lineage variation, and Y-linked gene content can be examined at chromosome scale for the first time in any canid.

### Per-Position Contiguity Check

To verify the gap-free property of Jindo1-G-TTAGGA at independently catalogued gap positions, we used the 13 internal gaps catalogued in a publicly available near-T2T canine reference^18^ (Beagle near-T2T 2025; LK_Cfam_Beagle_1.1; GCA_044048985.1) as reference coordinates. These gaps are distributed across nine chromosomes (per-gap coordinates in Supplementary Table S1). Per-position contiguity at all 13 coordinates was confirmed on both Jindo Hap1 and Hap2 by flanking-anchor mapping and whole-chromosome alignment (Methods). The detailed per-gap classification will be reported elsewhere.

### Assembly Quality, Completeness, and Repeat Annotation

Assembly polishing was performed in seven rounds following a strategy applied previously to other TTAGGA-grade assemblies. Rounds 1-3 applied structural variant (SV) polishing using Sniffles2^33^ /Jasmine^34^ and single-nucleotide variant (SNV) polishing using PEPPER-DeepVariant^35^ with trio-binned ONT reads. Rounds 4-6 repeated SV/SNV polishing with trio-binned HiFi reads. Round 7 applied an additional SV/SNV polishing cycle with merfin^36^ k-mer filtering. After polishing, local QV values were uniformly high (≥ 60) across all chromosomes except for a small number of terminal subtelomeric regions, which showed residual low-QV stretches. Zero-depth regions accounted for only 251,299 bp (0.011%) in Hap2 and 407,130 bp (0.017%) in Hap1.

Assembly completeness was evaluated using BUSCO v6.0.0 (carnivora_odb12) and the Genome Continuity Index (GCI). Jindo1-G-TTAGGA achieved BUSCO completeness of 99.3% (Hap1) and 96.4% (Hap2), and GCI of 98.2 (Hap1) and 94.7 (Hap2). Per-chromosome consensus QV after seven rounds of polishing is shown for both haplotypes in Figure 4A,B, and BUSCO completeness alongside publicly available canine reference assemblies is shown in Figure 4C. The Hap1 GCI value of 98.2 is in the range reported for current chromosome-scale references in other species. Repeat annotation was performed with RepeatMasker v4.2.3 against the Dfam 3.9 database, with total repeat content of approximately 32% on both haplotypes and LINE elements as the predominant class. This figure represents a lower bound, because canine-specific satellite families are not yet comprehensively represented in Dfam.

**Figure 4.**
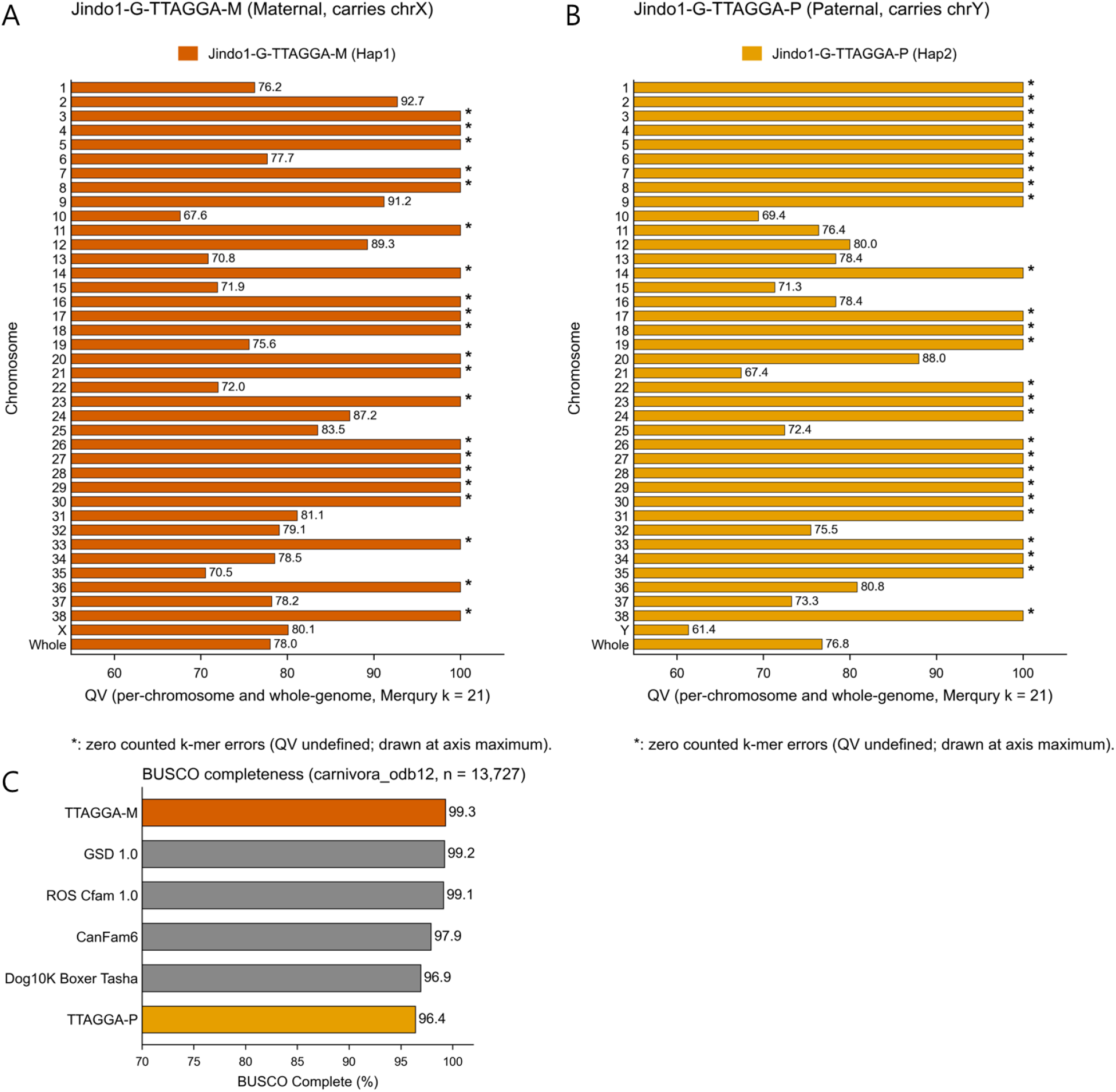
Assembly quality and completeness of Jindo1-G-TTAGGA. (A, B) Per-chromosome consensus QV after seven rounds of polishing for (A) Hap1 (Jindo1-G-TTAGGA-M; maternally inherited; chrX-carrying) and (B) Hap2 (Jindo1-G-TTAGGA-P; paternally inherited; chrY-carrying). Bars show Merqury QV (k = 21) for each of the 39 chromosomes and for the whole assembly (bottom row, “Whole”). Chromosomes with zero counted k-mer errors yield undefined (infinite) QV and are marked with an asterisk (*) at the axis maximum. (C) BUSCO completeness (carnivora_odb12, n = 13,727) for Jindo1-G-TTAGGA-M and Jindo1-G-TTAGGA-P compared with four publicly available canine reference assemblies (GSD 1.0, ROS_Cfam_1.0, CanFam6, and Dog10K Boxer Tasha).

## Discussion

The principal contribution of this work is a haplotype-resolved canine reference assembly in which every chromosome on both haplotypes is gap-free from telomere to telomere, including a single-contig 21.26 Mb Y chromosome. The assembly reaches zero internal gaps across both haplotypes, with a Hap1 GCI of 98.2 in the range reported for current chromosome-scale references in other species. The canine Y chromosome is, for the first time, available at a length and contiguity that allow direct comparison with the human and primate Y references that have driven sex-chromosome research over the past decade.

The Y chromosome is the most consequential extension of the present assembly. The 14 Mb of Jindo Hap2 chrY without counterpart in ROS_Cfam_1.0 raises the recovered canine chrY from approximately 15% at chromosome scale in ROS_Cfam_1.0 to approximately 79% of the cytogenetically estimated 27 Mb full-length chromosome in a single chromosome-scale contig. Although the inversion- and palindrome-rich proximal architecture suggested by the chrY synteny dotplot is informative, the assembly described here is primarily an enabling resource: ampliconic and palindromic region characterisation, identification of Y-linked gene families, paternal-lineage phylogenetics, and quantification of structural variation between male haplotypes are now possible at chromosome scale in the dog. Whether the additional 14 Mb is representative of canine chrY in general, or specific to East Asian breeds such as the Jindo, can be addressed only by comparable male assemblies from other lineages, a question this resource now makes tractable.

On chromosomes other than chrY, the assembly enables two observations that are themselves novel. The 8 Mb chr9 inversion signal observed in earlier canine-versus-canine alignments is shown to be an orientation artefact inherited in ROS_Cfam_1.0 rather than a Jindo-specific inversion. The Hap1 chrX is highly collinear with ROS_Cfam_1.0 chrX, with inverted regions accounting for less than 0.6% of total chrX length, indicating that the structural difference between male and female canine references in this assembly is essentially restricted to the Y chromosome.

We introduce TTAGGA to formalise a distinction that has become operationally relevant as long-read assemblies approach completeness: a chromosome may reach both telomeres yet retain internal gaps, a configuration observed in current near-T2T long-read assemblies. TTAGGA complements the T2T Consortium completeness criteria (Nurk et al. 2022) by making the zero-internal-gap requirement explicit, a property that is becoming a routine target for current long-read assemblies but is not separately named in existing terminology.

Several limitations of the present assembly warrant explicit statement. First, the recovered chrY (21.26 Mb) is approximately 6 Mb shorter than the cytogenetic estimate of ∼27 Mb. The unassembled fraction most likely corresponds to the heterochromatic p-arm satellite array and to ampliconic repeats whose tandem copy number exceeds the resolving power of current ONT ultra-long reads. Substantially higher coverage of reads longer than 100 kb, complementary technologies such as Strand-seq or optical mapping, or both, will likely be required to close it. Second, Hap2 chrY shows the lowest per-chromosome consensus QV in the assembly (QV 61.4, compared with the whole-haplotype QV of 76.8); the relative reduction reflects the difficulty of polishing dense repetitive content with long reads that map ambiguously, the same underlying cause of the residual low-QV stretches observed in subtelomeric regions of several other chromosomes. Third, the assembly is derived from a single male Jindo individual and is not a substitute for breed-level or species-level pangenomes; the Jindo population itself carries variation that is not represented here. Repeat annotation in canines also remains limited by the partial representation of canine-specific satellite families in Dfam, and the reported ∼32% repeat content is therefore a lower bound.

Two observations are relevant for subsequent canine assembly efforts. First, parental short-read data are required to label the two haplotypes as maternal or paternal, a label that, in any male assembly, is the basis for identifying the chrY-carrying haplotype. Reference-free phasing approaches (Hi-C, Strand-seq) can produce two haplotype-resolved assemblies but cannot make this assignment without trio data. Second, the combination of PacBio HiFi for initial diploid assembly and ONT ultra-long reads for gap bridging and chrY local assembly proved more effective than either platform alone. Future canine assemblies targeting the TTAGGA standard will likely require both platforms until single-platform performance at sufficient accuracy and read length becomes available.

The present work focuses on the assembly itself: its haplotype-resolved, gap-free, and chrY-inclusive properties, and the direct comparison with existing canine references. Variant discovery, repeat-class composition with canine-specific satellite annotation, detailed Y-chromosome architectural analyses, and population-level context for the Korean Jindo lineage will be reported separately.

## Materials and Methods

### Sample and DNA Extraction

High-molecular-weight genomic DNA was isolated from the whole blood of a healthy male Korean Jindo dog (Baeksan; Supplementary Figure S2) using the Monarch HMW DNA Extraction Kit for Tissue (T3060L, New England Biolabs), following the manufacturer’s protocol optimised for long-read sequencing. DNA quality and fragment size were assessed by pulsed-field gel electrophoresis and Qubit fluorometry, confirming that most fragments exceeded 100 kb.

### PacBio HiFi Sequencing and Read Processing

HiFi sequencing was performed on the PacBio Revio system across six flowcells, yielding 349.9 Gb (∼150×). Circular consensus sequencing reads were generated with ccs v6.4.0 using default parameters (mean QV > 29; mean read length 18-19 kb). Adapter sequences were removed with HiFiAdapterFilt^37^ . Reads were additionally trimmed 150 bp from both 5′ and 3′ ends to mitigate edge-quality degradation, then filtered to retain reads with length ≥ 1 kb and QV ≥ 20.

### Oxford Nanopore Sequencing and Read Processing

Ultra-long ONT reads were generated on the PromethION P2 Solo platform across 16 flowcells. Basecalling was performed using Dorado(v0.5.0)^38^. Reads were trimmed 2,000 bp from both ends to remove low-quality terminal bases, then filtered to retain reads with length ≥ 50 kb and QV ≥ 15, yielding∼76.8 Gb (∼30×; N50 101.3 kb).

### Illumina Trio Short-Read Sequencing and K-mer Analysis

Illumina short-read data were generated for both parents on the NovaSeq 6000 platform (the proband was not separately sequenced with Illumina short reads for trio binning, since trio binning uses only the parental k-mer databases). Adapter and low-quality base trimming was performed using fastp^39^ (front 10 bp, tail 5 bp; Q < 20 bases removed). Parental-specific k-mers (31-mer) were counted using yak with default parameters for trio binning. K-mer distributions were analysed with GenomeScope2 and Merqury (hapmer.sh, k = 31). Trio binning assigned the dam-derived haplotype to Hap1 (containing chrX in this male individual) and the sire-derived haplotype to Hap2 (containing chrY).

### Trio-binning and Multi-platform TTAGGA Assembly

Haplotype-resolved diploid assembly was performed with hifiasm v0.25.0-r726^30^ using the HiFi reads, ONT ultra-long reads, and parental yak k-mer databases (dual-scaf mode). The telomeric motif parameter was set to CCCTAA, which is the reverse complement of the canonical vertebrate telomeric repeat TTAGGG. Chromosome assignment used RagTag^40^ against ROS_Cfam_1.0 (Labrador Retriever, male). We use the term TTAGGA (Telomere-to-Telomere, Accurate, Gapless Genome Assembly) to refer to assemblies in which every chromosome (i) reaches both telomeres, (ii) contains zero internal gaps, and (iii) achieves a Merqury consensus QV of at least 60.

### Telomere Detection and ntLink Scaffolding

Telomeric repeats were identified using tidk^31^ with a threshold of > 100 repeats within 5,000 bp of scaffold ends. Additional scaffolding was performed with ntLink^41^ using ONT reads > 1 kb to join scaffolds with single-end or absent telomeres.

### Y-Chromosome Local Assembly

Sire-binned reads (HiFi and ONT, i.e., reads assigned to Hap2) were aligned to the dam-derived assembly (Hap1) using Winnowmap2 (meryl k-mer: 15 bp). Reads mapping to regions lacking telomeric ends were extracted (samtools −F 256) and assembled locally using hifiasm in haploid mode (n-hap = 1). The resulting assembly was resolved into a complete TTAGGA Y chromosome on Hap2 through iterative gap closing and polishing as described below. The final Hap2 chrY is a single 21,255,890 bp gap-free contig with TTAGGG telomeric repeats at the q-arm terminus.

### Gap Closing and Polishing

Gap closing was performed in ten rounds. Rounds 1-7 used a modified TGS-Gapcloser^42^ (without Racon polishing) with HiFi-based local assemblies^43^ and unplaced scaffold incorporation. Minimap2 alignment parameters for HiFi: −x map-hifi −P −c −s 100 −w 5 −U 10,1000000. Filtering criteria: aligned length > 500 bp, identity > 95%, unaligned length < 2,000 bp, alignment ratio > 95%. Rounds 8-10 employed manual scaffolding with large contig overlaps, trio-binned ONT Flye assemblies, and direct read-based gap filling (Minimap2: −x lr:hq; TGS-Gapcloser^42^: −−min_match 200, −−min_idy 0.97). Polishing was performed in seven rounds: Rounds 1-3 SV polishing with Sniffles2^33^/Jasmine^34^ and SNV polishing with PEPPER-DeepVariant^35^ on trio-binned ONT reads; Rounds 4-6 same pipeline with trio-binned HiFi reads; Round 7 additional SV/SNV polishing with merfin^36^ k-mer filtering. Low-QV regions were patched by identifying alternative high-QV contigs anchored to flanking assembly sequence.

### Assembly Quality Assessment

Assembly quality was assessed using: Merqury (QV and k-mer completeness; final Hap1 78.0, Hap2 76.8), BUSCO v6.0.0 (carnivora_odb12; Hap1 99.3%, Hap2 96.4%), Genome Continuity Index (GCI; Hap1 98.2,Hap2 94.7), and tidk for telomere end verification. Repeat annotation was performed with RepeatMasker v4.2.3 against the Dfam 3.9 database.

### Whole-Genome Synteny vs ROS_Cfam_1.0 (MUMmer / SyRI)

Whole-genome synteny was generated against ROS_Cfam_1.0 (Labrador Retriever; GCA_014441545.1) with MUMmer (nucmer + delta-filter) for both haplotypes, producing genome-wide alignment .delta and .coords files. SyRI^32^ was used to categorise structural variants. ROS_Cfam_1.0 was selected as the primary synteny reference because it provides the most contiguous chromosome-scale chrY among publicly available male canine references (3.94 Mb in a single chromosome-scale scaffold; see Table 3). Other male canine references (Wags, Yella v2, the Schall 2023 assemblies) recover chrY only as fragmented chromosome-scale and/or chrY-assigned unplaced scaffolds; female-derived references lack chrY entirely. The verified ROS_Cfam_1.0 chrY length (3.94 Mb; 3,937,623 bp; scaffold CM025139.1) was confirmed directly from the GCA_014441545.1 FASTA index.

### Per-Position Contiguity Check at Beagle near-T2T 2025 Internal Gap Positions

The Beagle near-T2T canine assembly^18^ was downloaded from NCBI. All internal N-stretches of length ≥100 bp on every chromosome were identified by sequential per-base scanning (Biopython SeqIO), recovering the 13 internal gaps reported in the deposited metadata, distributed across nine chromosomes. For each gap we extracted the ±5 kb non-N flanking sequences and aligned them separately to Jindo Hap1 and Hap2 using minimap2 (asm5 preset, −c −N 5). Whole-chromosome alignment between each gap-carrying chromosome and the corresponding Jindo Hap1/Hap2 chromosomes (minimap2 asm5 −c) was additionally used to confirm gap mapping. Detailed per-gap resolution classification and chr9 strand-orientation analysis will be reported elsewhere.

### Direct Measurement of chrY Length and Internal Gap Counts Across Public Canine References

All chrY lengths and internal gap counts reported in Tables 2 and 3 were measured directly from each reference FASTA by per-base scanning, rather than reproduced from literature or metadata. Internal gaps were defined as N stretches ≥ 100 bp at non-terminal positions (> 1 kb from scaffold ends). chrY-assigned scaffolds (chromosome-scale and unplaced) were identified from each assembly’s NCBI assembly_report.txt; chrY length was measured by inspecting the chromosome-scale sequence header and length, after manual verification that the largest sex-chromosome scaffold did not correspond to chrX. For Yella v2 (GCA_031165255.1), the NCBI assembly_report.txt does not explicitly assign chrY origin to any unplaced contigs in this assembly; we therefore tentatively assign chrY origin to the three largest unplaced contigs (DAOUOP010000040.1, DAOUOP010000041.1, DAOUOP010000042.1; 5.37 Mb total) based on their large size and the absence of a chromosome-scale chrY scaffold in this assembly. The eight publicly available male canine assemblies surveyed are: ROS_Cfam_1.0 (Labrador Retriever, isolate SID07034; GCA_014441545.1)^15^, Wags^12^(the male Basenji assembly; GCA_004886185.2), Cairn Terrier CA611 (GCA_031010295.1) and Bernese Mountain Dog OD (GCA_031010635.1)^13^, Yella v2 (Labrador Retriever; GCA_031165255.1)^14^, Yella alternate haplotype (GCA_012044875.1), Yella principal haplotype (GCA_012045015.1), and the Whippet 2024 PacBio HiFi Revio assembly (GCA_043643935.1). GenBank accessions and per-assembly chrY measurements are tabulated in Supplementary Table S2.

## Data Availability

All raw sequencing reads and the Jindo1-G-TTAGGA assembly will be deposited in public repositories; accession numbers will be provided upon publication of the accompanying full-length manuscript. Assembly data are also available at https://kogic.kr/Dognomics. Detailed supplementary tables (S1-S5) are available upon request and will be deposited together with the accompanying full-length manuscript. For collaboration access, please contact the corresponding author at.

## Author Contributions

J.B. (Jong Bhak) and S.L. (Sung-kyung Lee) supervised the study and designed and outlined the project. Y.K. (Yoonsung Kwon) performed the telomere-to-telomere genome assembly. J.K. (Jong-Seok Kim) was responsible for overall sample collection and management. H.C. (Hyoungjin Choi) performed the bioinformatic analyses and wrote the manuscript. S.P. (Sangsoo Park) generated all sequencing libraries and performed sequencing. Y.C. (Yookyung Choi) performed the mitochondrial genome assembly. J.B. (Jihun Bhak), D.S. (Donghyun Shin), and K.A. (Kyungwhan An) reviewed the manuscript. D.R. (Doug-Young Ryu), W.P. (Woon Kee Paek), D.P. (Daeui Park), S.J. (Sungwon Jeon), J.M.K. (Jaemin Kim), M.H.S.S. (Mikkel-Holger S. Sinding), Y.C. (Youngjae Choe), and B.H. (Bo-Ra Hyun) provided scientific advice and reviewed the manuscript.

### Acknowledgments

We thank the Korean Jindo and Domestic Animals Center (Jindo, 58915, Republic of Korea) and the Korea Heritage Service (Daejeon, 35208, Republic of Korea) for their support. We thank Ulsan City and UIPA for supporting various genome projects in UNIST.

