## Supplementary Figures S1-S2 and Tables S1-S5 for "Telomere-to-telomere, accurate, and gapless genome assembly (TTAGGA) of the Korean Jindo dog with a single-contig Y chromosome"

### Supplementary Materials

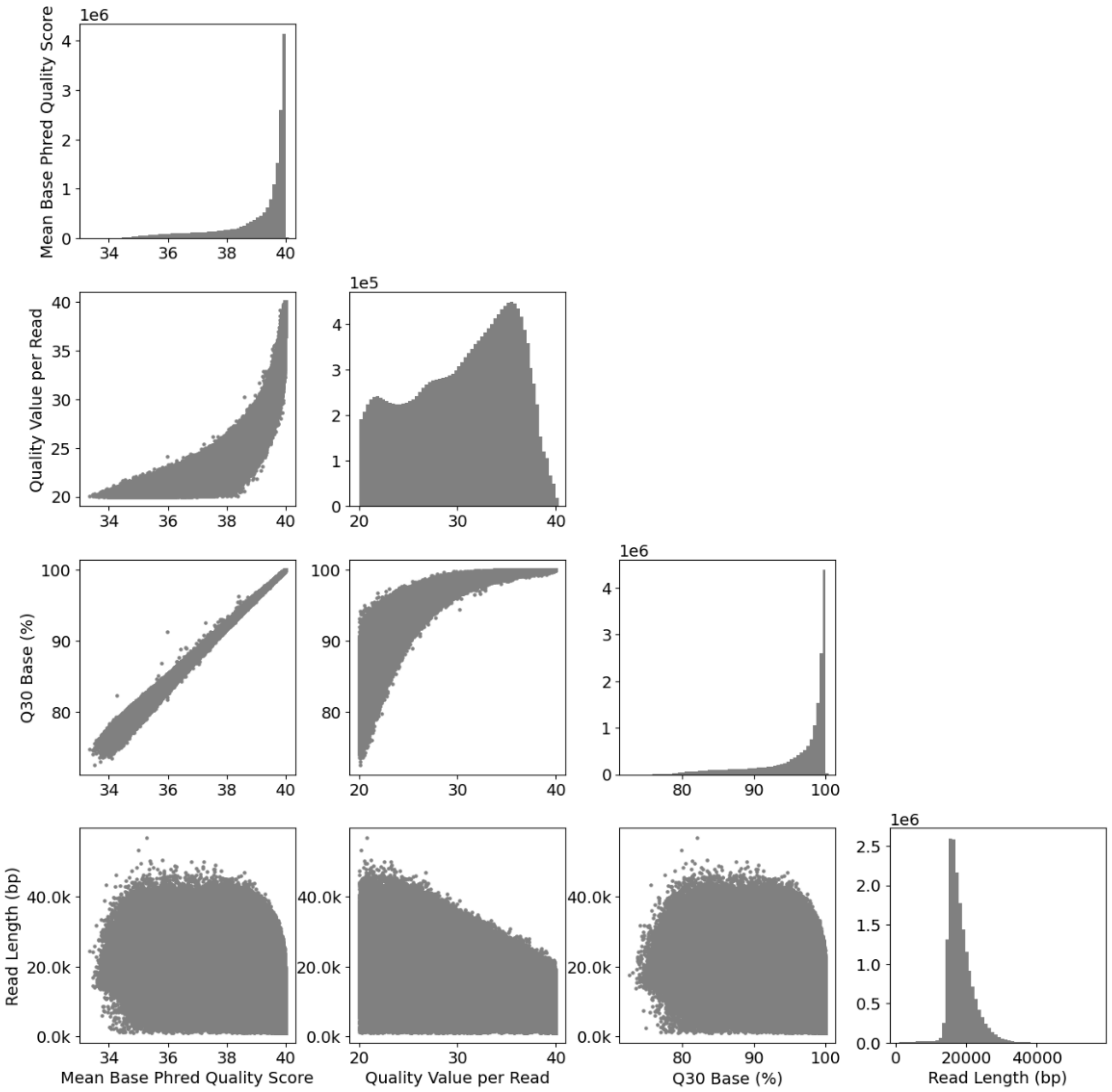


**Supplementary Figure S1a. Quality metrics of PacBio HiFi sequencing data across six Revio flowcells** Scatter-matrix display of per-read mean Phred quality score, quality value per read, Q30 base fraction, and read length distribution. After trimming 150 bp from both ends and filtering (length ≥ 1 kb; QV ≥ 20), a total of 317.6 Gb (17.3 million reads) were retained for assembly.

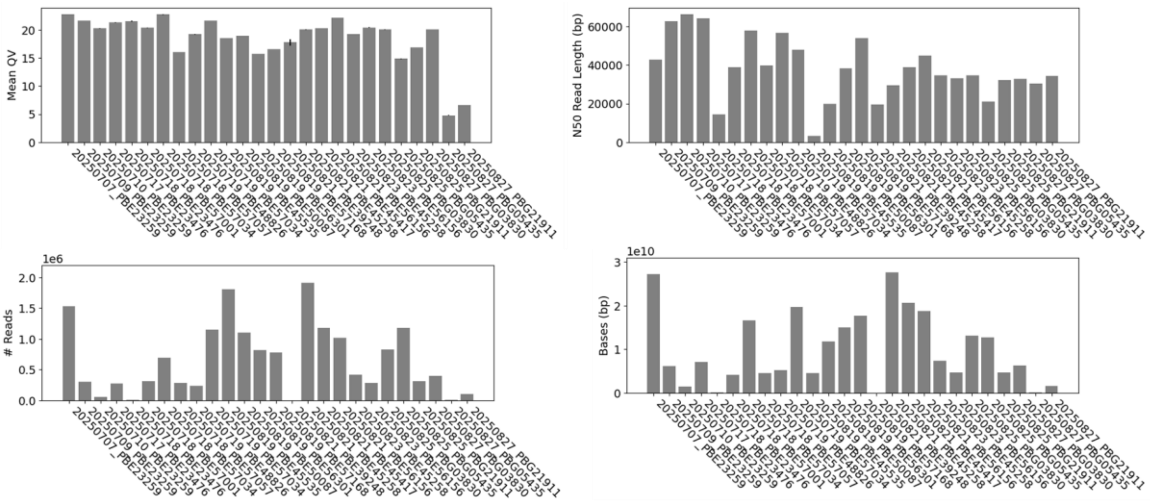


**Supplementary Figure S1b. Quality metrics of Oxford Nanopore ultra-long read data across 16 PromethION P2 Solo flowcells** Per-flowcell mean QV, N50 read length, read count, and total yield (bases). The combined dataset yielded 258.8 Gb with a total N50 of 37.8 kb. Ultra-long reads (length > 50 kb; QV > 15) accounted for ~85.1 Gb (~34×). After trimming 2,000 bp from both ends, 76.8 Gb (~30×; N50 101.3 kb) were retained.

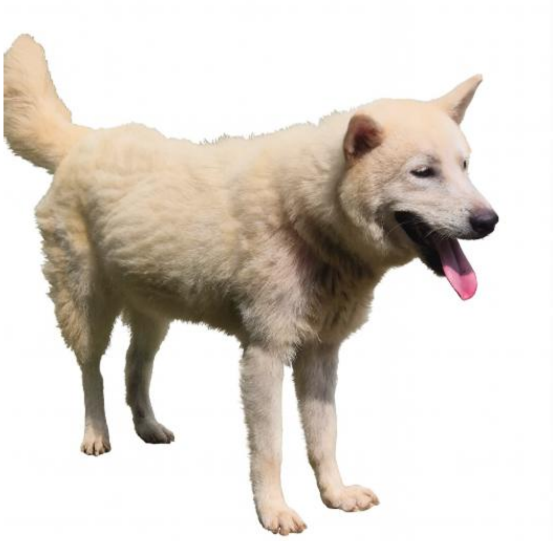

**Supplementary Figure S2. Baeksan, a purebred Jindo dog under official breed management in Korea** Baeksan is a purebred Korean Jindo dog (designated as Korean Natural Monument No. 53), a breed that is nationally protected and systematically managed in Korea for maintaining its genetic integrity. As a representative individual of this indigenous and genetically distinct East Asian lineage, Baeksan was selected for generating the Jindo1-G-TTAGGA reference genome.

- Supplementary Table S1. Per-gap coordinates and 5 kb flanking-anchor classification of the 13 internal gaps in the Beagle near-T2T 2025 assembly against Jindo1-G-TTAGGA Hap1 and Hap2.
- Supplementary Table S2. GenBank accession numbers and per-assembly chrY measurements for the eight publicly available male canine genome assemblies surveyed in Table 3.

- Supplementary Table S3. Per-haplotype assembly statistics (scaffolds, scaffold/contig N50, largest/smallest scaffold, GC content, Merqury QV, switch error).
- Supplementary Table S4. RepeatMasker repeat-class composition for Hap1 and Hap2.
- Supplementary Table S5. Full SyRI structural-variant categorisation between Jindo1-G-TTAGGA (Hap1, Hap2) and ROS_Cfam_1.0 across all variant classes (syntenic, inversion, translocation, duplication, copy gain/loss, highly diverged regions, SNPs, insertions, deletions).
